# LITOS - a versatile LED illumination tool for optogenetic stimulation

**DOI:** 10.1101/2022.03.02.482623

**Authors:** Thomas Christoph Höhener, Alex Landolt, Coralie Dessauges, Paolo Armando Gagliardi, Olivier Pertz

**Affiliations:** Institute of Cell Biology, University of Bern, 3012 Bern, Switzerland; ETH Zurich, Department of Biosystems Science and Engineering, 4058 Basel, Switzerland

## Abstract

Optogenetics has become a key tool to manipulate biological processes with high spatio-temporal resolution. Recently, a number of commercial and open-source multi-well illumination devices have been developed to provide throughput in optogenetics experiments. However, available commercial devices remain expensive and lack flexibility, while open-source solutions require programming knowledge and/or include complex assembly processes. We present a LED Illumination Tool for Optogenetic Stimulation (LITOS) based on an assembled printed circuit board controlling a commercially available 32×64 LED matrix as illumination source. LITOS can be quickly assembled without any soldering, and includes an easy-to-use interface, accessible via a website hosted on the device itself. Complex light stimulation patterns can easily be programmed without coding expertise. LITOS can be used with different formats of multi-well plates, petri dishes, and flasks. We validated LITOS by measuring the activity of the MAPK/ERK signaling pathway in response to different dynamic light stimulation regimes using FGFR1 and Raf optogenetic actuators. LITOS can uniformly stimulate all the cells in a well and allows for flexible temporal stimulation schemes. LITOS’s ease of use aims at democratizing optogenetics in any laboratory.

## Introduction

While optogenetics was first used in neurobiology^1^, it is now widely used to control a wide variety of cell biological processes with high spatio-temporal resolution^2^. This has been made possible by the discovery of a number of light-responsive protein domains that were engineered to build actuators to control almost any cell biological process^2^. The precision of optogenetics to perturb cellular systems can lead to a deeper understanding of their dynamic regulation^3^. However, optogenetic experiments also require appropriate optical stimulation hardware.

A classical optogenetic experimental setup is to use automated light microscopes to stimulate cells expressing optogenetic actuators and record any desired cellular output. When combined with spectrally compatible biosensors in a fluorescence microscope, this setup can both control the cellular input using the optogenetic actuator and record the output dynamics using the biosensor^4^. This has proven a very powerful approach to study signaling dynamics^5–7^. Unfortunately, the field-of-view of the microscope’s lens system limits the number of cells that receive the light stimulation. Thus, microscopes cannot stimulate sufficient amounts of cells to measure cellular outputs using biochemical methods. Further, the number of different input stimulation patterns that can be induced in parallel is restricted. Finally, any long-term optogenetic control of cells on time scales of multiple days might be impractical or too expensive, especially in microscopy facilities.

Using a dedicated illumination source to separate the stimulation from the imaging process circumvents some of these limitations. LED stripes mounted in an incubator can be used to stimulate an optogenetic actuator in a large number of cells, opening up the possibility to measure cellular outputs using biochemical methods such as western blot, proteomics or transcriptomics^8^.

More advanced setups combine microcontrollers and light sources to specifically illuminate individual wells from multi-well plates. This allows to stimulate multiple wells with different patterns of light inputs in parallel, resulting in higher experimental throughput. A number of hardware solutions that take advantage of LEDs as illumination sources have been developed^9–12^. Those open-source devices are fairly cheap, but require programming knowledge and might rely on manufacturing expertise not available in most laboratories. Recently, some commercial products (e.g. LUMOS from AXION Biosystems) have entered the market, but their cost remains high. A cheap, easy-to-assemble, user-friendly device that provides experimental flexibility is still missing in the optogenetic community.

Here we present a new LED Illumination Tool for Optogenetic Stimulation (LITOS) to democratize optogenetics in any cell biology laboratory. LITOS allows high throughput dynamic light stimulation of large cell populations in different multi-well plate formats, petri dishes or cell culture flasks. Its user-friendliness allows users with little technical knowledge and no coding skills to easily use the device and set up complex illumination patterns. Moreover, LITOS can be assembled without soldering and does not require 3D printing of any component. We provide a detailed description of LITOS assembly, and we document a user-friendly GUI solution to configure complex illumination patterns. We validated LITOS using mammalian cell lines expressing optogenetic actuators based on FGFR1 and Raf, and evaluating the mitogen-activated protein kinases (MAPK)/extracellular signal-regulated kinase (ERK) pathway as a signaling output, using both biochemical and imaging approaches.

## Results

### An easy-to-assemble LED illumination device

To minimize the need for complex manufacturing procedures, we based LITOS entirely on commercially available components (Fig. 1 and Fig. 2a). LITOS relies on a custom printed circuit board (PCB), which can be ordered pre-assembled from a PCB manufacturer. The PCB acts as a control unit for a commercially available RGB LED Matrix. The PCB contains an ESP32 microcontroller, which provides sufficient computational power to control the LED matrix and a wireless network node (Fig. 2b). The RGB LED matrix consists of an array of 32×64 LEDs with a 3 mm pitch that allows for illumination with red, green and/or blue light. This can illuminate cells cultured in 6, 12, 24, 48 or 96 multi-well plates, as well as petri-dishes and cell culture flasks (Fig. 2c). We choose this 3 mm pitch LED matrix format because it perfectly aligns with the typical 96 well plate (9 mm) layout, resulting in a 2×2 LED array centered in each well. In addition, the RGB LED matrix allows the user to modulate the intensity of the different colors at will. The PCB module and the RGB LED matrix are connected with a flat ribbon cable and a power cable. This simple assembly procedure does not require any soldering. The schematics of LITOS are described on our GitHub page (https://github.com/pertzlab/LITOS).

**Figure 1:**
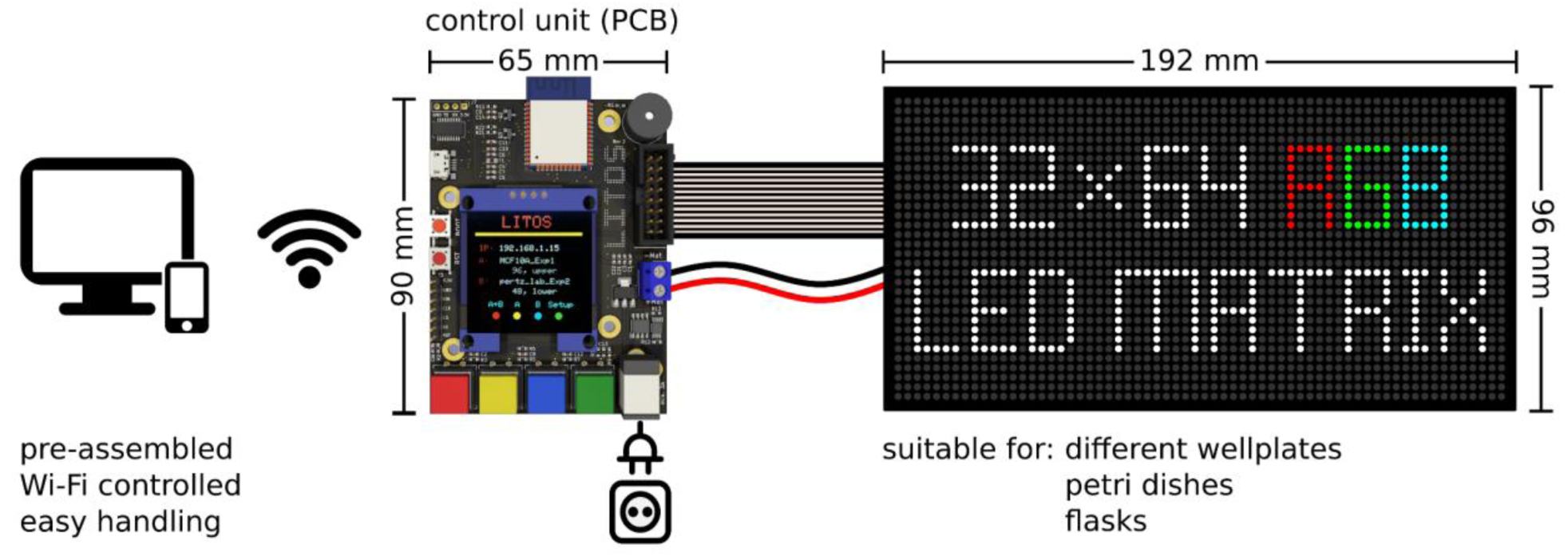
Schematics of LITOS components. LITOS is composed of two parts: a printed circuit board (PCB) which acts as a control unit and a commercially available 32 × 64 RGB LED matrix. LITOS can be controlled with a computer or smartphone over a WLAN connection. The PCB controls the 32 × 64 RGB LED matrix to illuminate optogenetic cells. The densely pitched LED matrix enables the illumination of various cell culture dishes and well plates. With a screen and four buttons, LITOS can be directly controlled and can display messages regarding the execution of an experiment or about actions that require user intervention.

**Figure 2:**
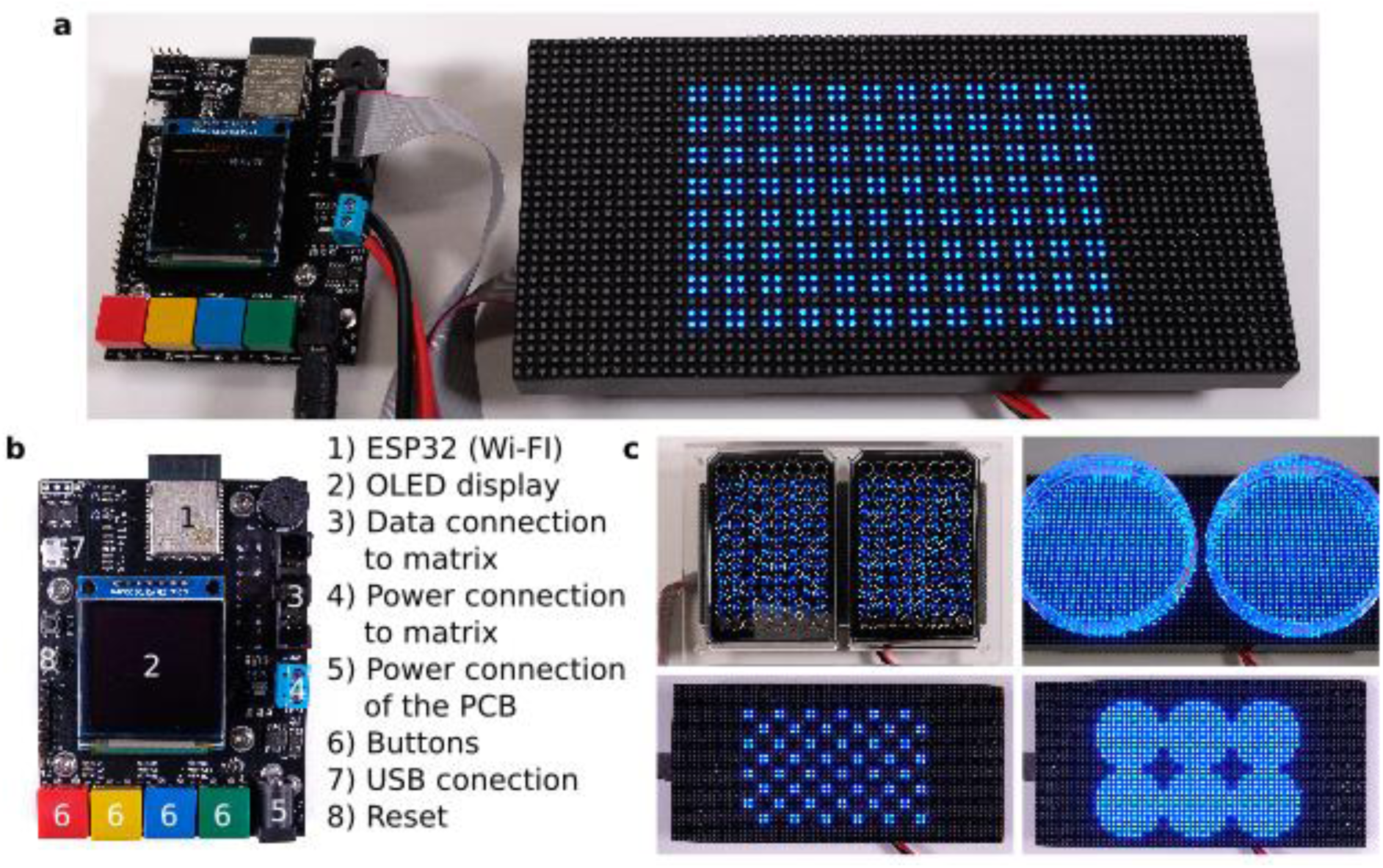
LITOS can generate flexible light stimulation patterns. a) Upon assembly, LITOS is composed of the PCB and 32 × 64 LED matrix. Here, LITOS is used to generate an 8 × 12 matrix of 4 LEDs per well to stimulate each well of a 96 well plate. b) The PCB provides everything needed to operate the matrix: the microcontroller with WLAN module, the connections to the LED matrix, a display and four buttons for user-defined actions (e.g. marking a plate outline, experiment start and experiment termination). c) Examples of LITOS applications with different plates or dishes formats. Left panel: simultaneous stimulation of two 96 well plates, using an optional plexiglass mask for plate alignment. Top right: two 100 mm petri dishes. Bottom left: alternated illumination pattern for a single 96 well plate. Bottom right: stimulation pattern for a 6 well plate.

To easily align the plates in dark environments, we implemented 3 solutions: i) using LITOS to display a reference outline of the desired well plate on the LED matrix, which can be directly activated by a button on the PCB (Supplementary Fig. 1a); ii) aligning the plate to the upper left corner of the LED matrix (Supplementary Fig. 1b) and iii) using a mask that positions the plate on the LED matrix (the latter can be produced by 3D printing or CNC milling) (Supplementary Fig. 1c). The latter solution also provides the possibility to place and stimulate two multi-well plates on one LITOS device simultaneously.

The overall cost of LITOS, which comprises a custom PCB assembled by a PCB manufacturer, and an off-the-shelf LED matrix, accounts for approximately USD 150.00. This low price of the device allows for easily scaling up experiments by ordering more units. In this case, the price per unit can even further decrease.

### User-friendly software for facile set up of optogenetic experiments

As for the hardware, LITOS’s software was designed to be user-friendly. The software present on the control unit uses user-defined illumination schemes, which are then processed to control the LED matrix (Fig. 3a-c).

**Figure 3:**
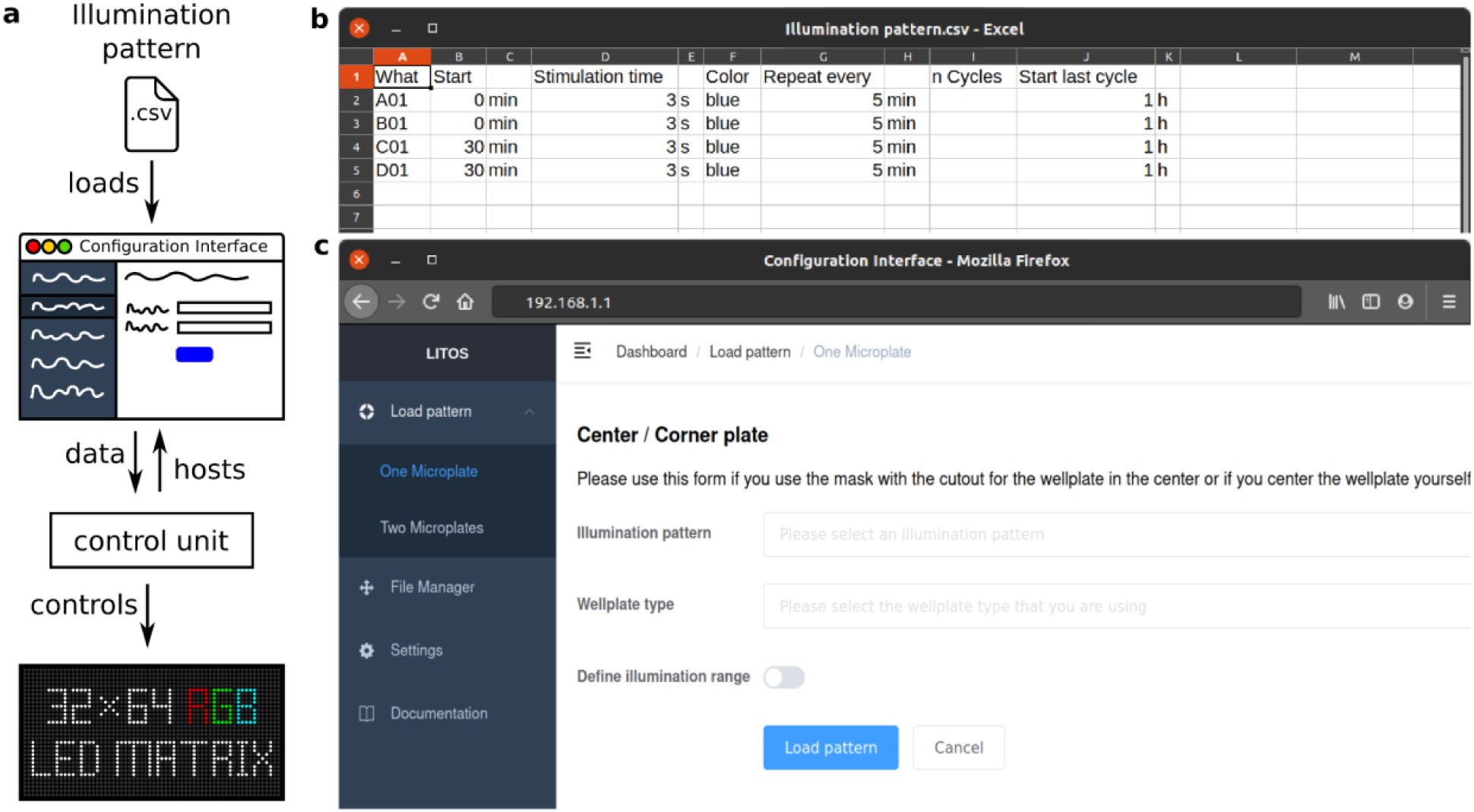
LITOS general user workflow. a) Schematic of the experimental workflow to generate specific illumination patterns. A csv file with the desired pattern can be uploaded via a web browser to the “Configuration Interface” hosted by the control unit. The control unit then operates the LED matrix according to the uploaded illumination pattern. b) Example csv file used for a specific illumination pattern. Here, wells A01 and B01 are programmed to be illuminated every 5 min with a 3 s pulse of blue light; Wells C01 and D01 are programmed with the same pattern but with a delay of 30 min. In both cases the experiment has a total duration of 1h. c) The User-interface hosted by LITOS is used to transfer the illumination patterns, which can subsequently be loaded for optogenetics experiments.

No knowledge of programming is needed to set up desired illumination patterns. To define which LEDs should be switched on, at which specific time, for which specific duration, and in which specific color, the user creates a comma separated value file (.csv). This can be done using spreadsheet applications, such as Google Sheets, Microsoft Excel or LibreOffice Calc. In the csv file, each line describes an illumination command (Fig. 3b). Each column indicates different parameters of the stimulation program: LEDs to be activated, starting time of illumination, illumination duration, color and intensity of the active LEDs. Other columns allow the generation of loops in case the illumination should be repeated several times (e.g. illuminated every 10 min for 3 hours). So that the user does not always have to define individual LEDs, we designed a number of keywords for: i) Well (e.g. “A02” / “A2” for Well A02); ii) Column and row (e.g. “D” for the whole row D); iii) whole multi-well plate (Plate); iv) whole matrix (Whole); v) circle and rectangles (Rec_start_end). LITOS’s software translates these keywords into the corresponding LED positions of the matrix.

Additionally, within such csv files, custom messages with countdowns can be programmed to be displayed on the control unit. This can help the user with any additional tasks during the optogenetic experiment, such as pipetting a reagent to a specific well, for example. As these tasks can be time-sensitive, a small piezo buzzer supports the user with additional auditive signals beside the visual ones. The csv file of the illumination pattern is then transferred to the ESP32 microcontroller via its Wi-Fi module. A java-script based configuration interface (Fig. 3c) is hosted on the control unit to upload and manage illumination patterns, change settings and prepare LITOS for an experiment. Multiple illumination patterns can be stored on the control unit’s internal memory, allowing for reusing stimulation patterns without the necessity of re-uploading them. The control unit’s configuration interface can either be accessed over a WLAN access point created by LITOS or over the local Wi-Fi network. The information needed for the connection, such as LITOS’s IP address or the name of the created WLAN access point, are displayed on the internal display. When connected, the user can access the configuration interface by a web browser on any computer or smartphone independently of the platform.

Once an illuminating pattern is transferred and selected, LITOS is ready to start the experiment. The four buttons present on the control unit allow the user to directly start or abort the experiment, or to indicate with an outline on the RGB matrix where to place the multi-well plate. Execution of illumination programs on LITOS can be directly monitored on the control unit’s display. Supplementary Movie S1 shows the LED matrix applying a stimulation pattern in which every column of a 96 well plate is illuminated sequentially.

While performing initial experiments, we found that the LED matrix developed heat that possibly could be detrimental to cell health. We found that the LED matrix heats up even when idle, i.e. when no light stimulation occurs. To solve this issue, we created a circuit using N- and P-metal–oxide–semiconductor field-effect transistors (MOSFETs) to deliver electric current to the matrix only when an illumination process is required. This reduced the heat development observed for repetitive stimulation patterns. We then evaluated the change in medium temperature due to LITOS illumination by measuring the medium temperature after different stimulation patterns. As expected, our measures show that longer and more frequent light stimulation led to higher temperature increase, which reached a plateau after around 1 hour (Supplementary Fig. S1). However, the experiments shown in this article never required light stimulation schemes that would produce enough heat to compromise cell health. In case experiments require very sustained light inputs, adding small passive heat sinks might allow to reduce overheating. Another approach is to engineer the optogenetic module with a low k_off_ to cause sustained light-dependent interactions. For example, in the iLID (improved light inducible dimer) system, the affinities and lifetimes of the light-dependent interaction can be engineered by diverse mutations^13^.

### Showcasing LITOS in the study of MAPK/ERK signaling dynamics

LITOS lends itself to a wide spectrum of optogenetic applications and model systems. To demonstrate the flexibility and versatility of LITOS, we performed optogenetic experiments to study the MAPK/ERK pathway. The MAPK network is able to decode a variety of extracellular and intracellular cues, to convert them into specific fate decisions, and plays pivotal roles in both physiology and pathology^14^. For this reason, multiple optogenetic actuators and fluorescent biosensors have been developed to both control and measure different signaling nodes of this pathway, making it an ideal model system to showcase LITOS.

We used mouse NIH3T3 fibroblasts and MCF10A mammary epithelial cell lines, engineered to stably express a genetically-encoded circuit consisting of an optogenetic actuator to activate different nodes in the MAPK network, and a fluorescent biosensor to record the dynamics of the ERK signaling output. This circuit consists of an optogenetic fibroblast growth factor receptor (optoFGFR), that consists of a myristoylated intracellular domain of the fibroblast growth factor receptor 1 fused with the light sensitive domain CRY2 that upon stimulation with blue light multimerizes, autophosphorylates, and activates the MAPK pathway^4^. The ERK signaling output can then be measured using an ERK kinase translocation reporter (KTR) that reports on ERK activity at single-cell resolution^15^. To be spectrally compatible with optogenetic activation, ERK-KTR is fused with a mRuby2 red fluorescent protein that can be excited without activating optoFGFR. In addition, both cell lines stably express a histone H2B nuclear marker fused to the far red fluorescent protein miRFP703 (H2B-miRFP703). A scheme of this genetic circuit is shown in Figure 4a. We then used a computer vision approach to infer ERK activation status for each cell (Fig. 4b). For that purpose, nuclei were automatically segmented using the H2B-miRFP703 channel and used to derive a nuclear mask and a cytosolic ring mask around the nucleus. Then, a ratio of the median fluorescence intensities of the ERK-KTR-mRuby2 channel using the cytosolic and nuclear masks is calculated as a proxy for ERK activity (C/N ratio).

**Figure 4:**
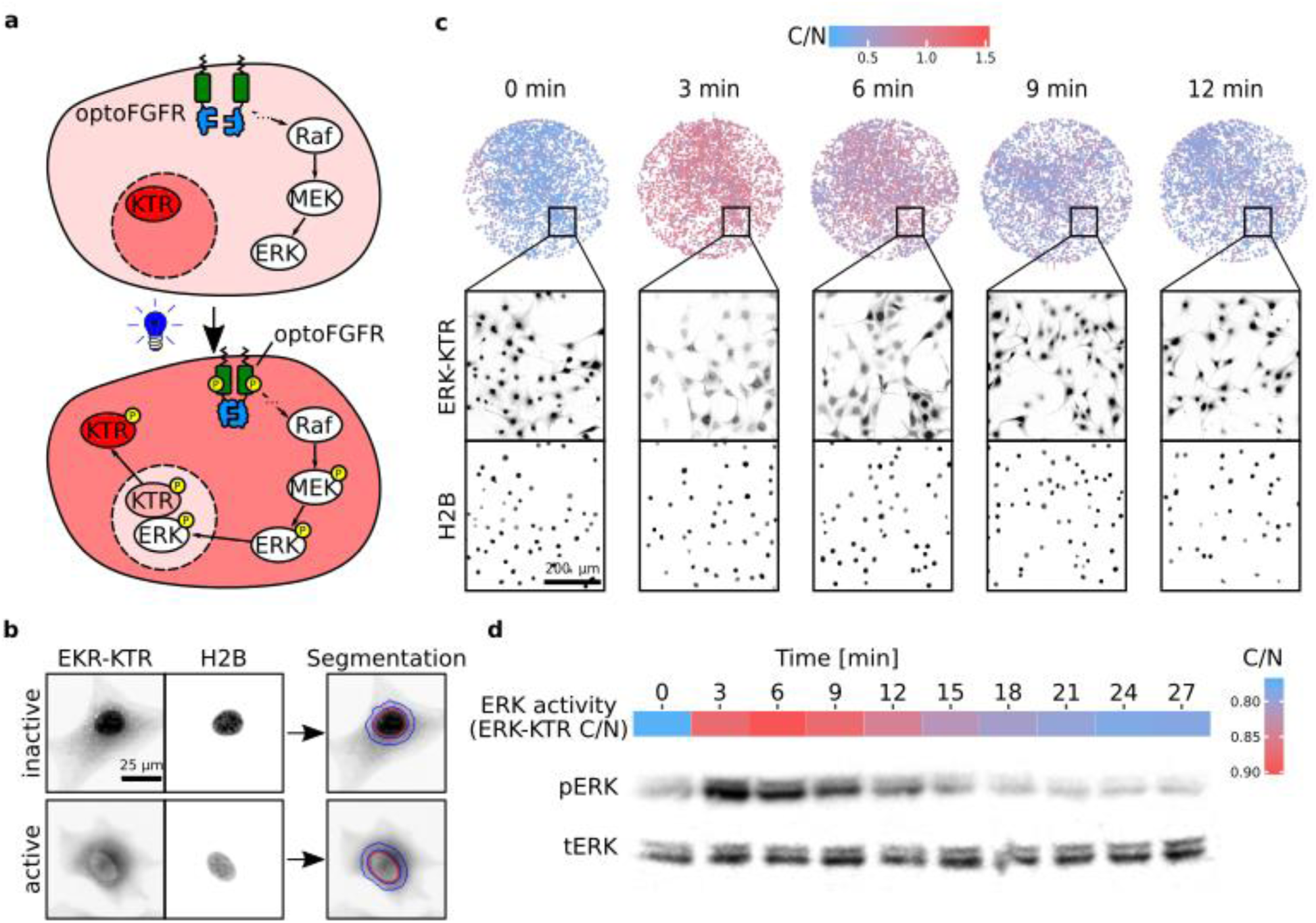
LITOS in the study of MAPK pathway signaling dynamics. a) Upon stimulation of optoFGFR, the MAPK pathway gets activated, leading to the phosphorylation and subsequent activation of ERK. Active ERK phosphorylates ERK-KTR, leading to its reversible translocation into the cytosol. b) Computer vision pipeline for image analysis. Nuclear segmentation of NIH3T3 cells (red), based on the nuclear marker H2B, and its ring-shaped expansion (blue) are used to measure the ratio between cytosol and the nucleus (C/N) of ERK-KTR. High C/N ratio corresponds to high ERK activity. c) The upper row represents the color-coded ERK-KTR C/N ratio of all NIH3T3 cells in 6.38 mm diameter wells (96 well plate). The two lower rows represent a full resolution example of a FOV that composes the larger image above, with H2B-miRFP703 and ERK-KTR-mRuby2 shown in inverted grayscale. d) MCF10A cells expressing ERK-KTR were stimulated with a 10 s long light pulse, fixed with 4 % paraformaldehyde and their ERK activity was quantified as shown in b. The ERK activity was then compared to a western blot against phosphorylated and total ERK from the same cells (below). Native, non-cropped western blots are available in Supplementary Figure S2.

The first feature of LITOS that we evaluated was the ability of the LED array to homogeneously illuminate all the cells within one well. This feature is crucial for whole-population applications. For this purpose, we stimulated 96-well plates, in which each well is illuminated by four LEDs, and measured ERK-KTR C/N in all the cells of a well by acquiring a large mosaic picture of each entire well. Displaying the ERK-KTR C/N ratio values of each single cell in entire wells revealed that the four LEDs homogeneously illuminate a single well, inducing robust optogenetic activation of MAPK pathway (Fig. 4c). Because the different experiments were performed in adjacent wells, this also shows that there is no light spillover from one well and another.

We then tested LITOS’s capability to generate robust ERK activation detected by population average biochemical methods such as western blot. To produce sufficient protein lysate we seeded MCF10A cells in 6 well plates, yielding about 250 µg of protein lysate per well. Illumination of an entire well of a 6-well plate required switching on 112 LEDs (Fig. 2c, lower right panel). We stimulated the optoFGFR-expressing cells using a 10-second pulse of blue light with LITOS and fixed the cells using paraformaldehyde at different time points with a 3-minute interval. To compare the ERK activation dynamics observed using western blot using a phosphoERK antibody (pERK) versus the ERK-KTR biosensor, we measured ERK-KTR C/N ratio in the same wells prior to protein extraction. To average any local variability of single-cell ERK-KTR C/N ratios, we sampled each well with 16 FOVs and computed their population average. The western blot measurement of endogenous pERK closely follows the measurement of ERK kinase activity using the ERK-KTR biosensor. Consistently with previous observations^5,16^, both pERK and ERK-KTR signals displayed a steep increase in ERK activity directly after light stimulation, followed by slower adaptation, that lead to pre-stimulation ERK activity levels in appropriately 20 minutes. The ERK-KTR signal (peak at 6 min) was slightly delayed compared to the pERK signal (peak at 3 min) (Fig. 4d). This might reflect the time that ERK needs to translocate to the nucleus and/or the time that the ERK-KTR biosensor needs to translocate to the cytosol.

These experiments show that LITOS can uniformly and reliably activate all the cells in a large well, demonstrating that it is suitable for stimulation of large cell numbers required for western blot, making it also compatible with other population-average measurements (e.g. transcriptomics or (phospho)proteomics).

### LITOS can produce simple as well as complex dynamic stimulation patterns

To fully exploit the potential of optogenetics in controlling dynamic biological processes, LITOS should be capable of activating different optogenetic actuators, as well as having high flexibility in creating complex illumination patterns. We therefore performed a series of additional experiments to demonstrate that LITOS meets these requirements. In all the following experiments, we programmed LITOS to automatically apply the light inputs to different wells at specific times so that the cells in a 96-well plate can be fixed simultaneously at the end of the experiment. Upon cell fixation, we then imaged all the wells of the plates using automated microscopy. We then took advantage of our automated image analysis pipeline ^16^ to calculate a mean ERK activity per well. Note that performing such experiments with pipetting of growth factors to activate the MAPK network in a time-resolved manner would be highly challenging from a logistics perspective.

We first evaluated if LITOS is also able to stimulate an optogenetic actuator that uses a different activation mechanism than the one of optoFGFR. Contrary to optoFGFR, which is based on the multimerization of a light-sensitive membrane receptor, optoRaf^17^ involves light-dependent recruitment of a catalytic c-Raf domain to the membrane through a CRY2 system. To evaluate if LITOS is also able to stimulate optoRaf, we screened light inputs of different durations, and compared them to optoFGFR stimulation. We observed that a 0.5 minutes light pulse induced higher ERK amplitude in the optoRaf versus the optoFGFR system (Fig. 5a). This was also apparent at later time points. In response to increasing light input, optoFGFR led to a transition from transient to sustained ERK dynamics, as previously observed^5^, while optoRaf always displayed adaptive ERK dynamics. The different ERK dynamics induced by optoFGFR and optoRaf inputs might reflect different mechanisms of actuation. Alternatively, it could reflect a difference in the ability of the two actuators to activate ERK from different levels within the MAPK network: optoFGFR triggers the activation of a full MAPK network downstream of the receptor, whereas optoRaf only triggers the activation of an isolated MEK/ERK network. This illustrates how LITOS can be used to study ERK dynamics induced by different optogenetic actuators.

**Figure 5:**
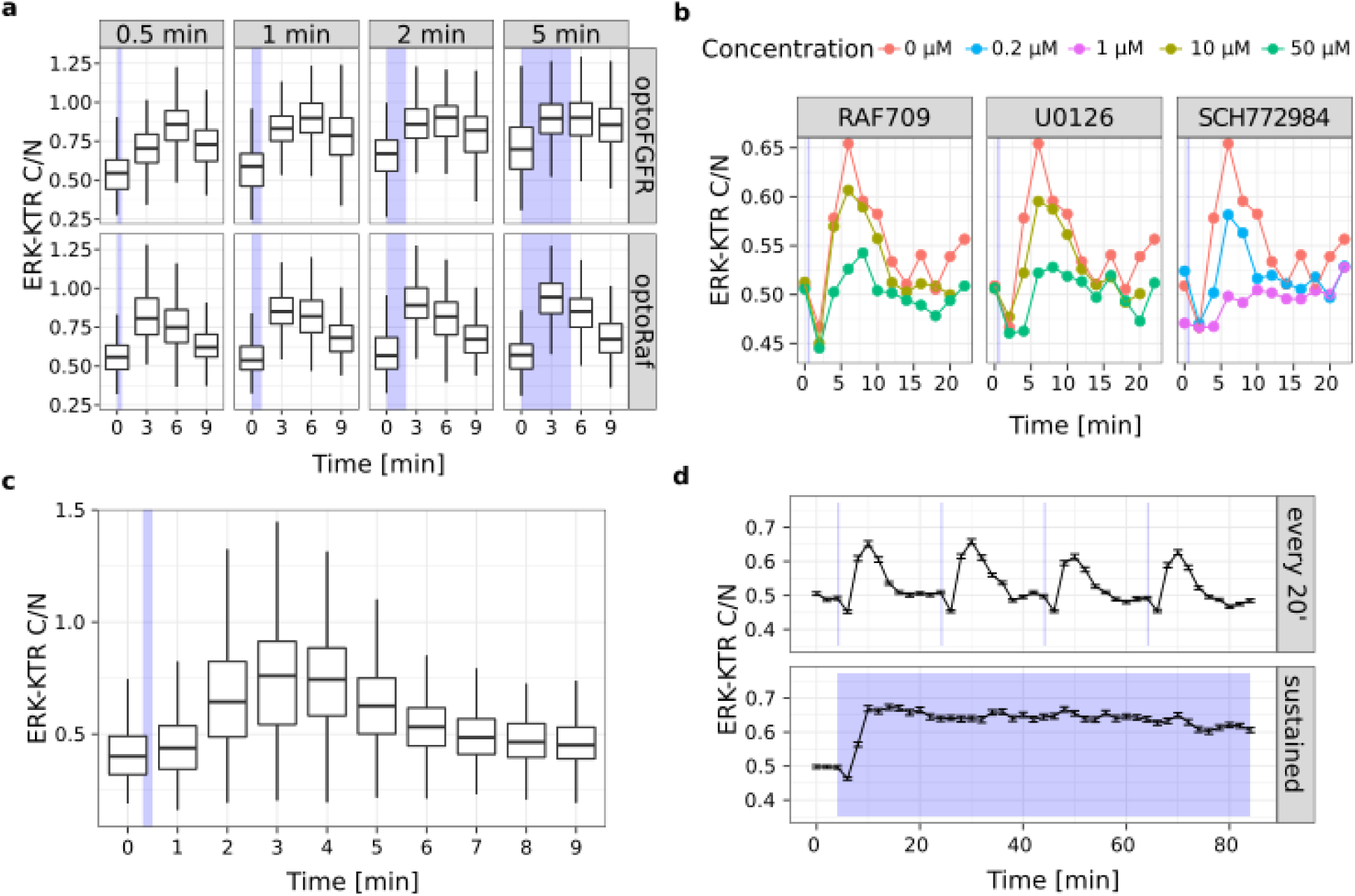
Showcasing LITOS’s versatility in the study of MAPK signaling dynamics. a) ERK response to optoFGFR or optoRaf light inputs of different times. b) Small drug screen on NHI3T3 expressing optoFGFR performed in a single 96 well plate using different concentrations of the Raf inhibitor RAF709, the ERK inhibitor SCH772984 and the MEK inhibitor U0126. c) ERK activity dynamics measured after a single 2 s long blue light pulse with LITOS measured with a temporal resolution of 1 min. d) Comparison of pulsatile and sustained ERK activity dynamics generated with LITOS. a-d) ERK activity was measured as ERK-KTR C/N. Box plots represent the median (central line), 2nd and 3rd quartile (boxes) and 1.5 interquartile range (whiskers) of single-cell ERK-KTR C/N measurements. Line plots represent the average (line) and standard error of mean interval (error bars) of single-cell ERK-KTR C/N measurements.

We then further evaluated the potential of LITOS to perform a drug screen with ERK dynamics as a readout (Fig. 5b). As proof of concept, we tested 3 drugs that inhibit the MAPK pathway: the Raf inhibitor RAF709, the MEK inhibitor U0126 and the ERK inhibitor SCH772984. We applied each inhibitor at 2 concentrations. We stimulated optoFGFR cells with a single 10 seconds blue-light pulse in presence of the inhibitors, and measured ERK activity at 12 different time points with a 2 minutes time resolution. All three drugs led to a dose-dependent reduction of ERK activity. The peak of ERK activity was markedly reduced with the two highest concentrations of RAF709 and U0126, or even abrogated with the highest concentration of SCH772984 (Fig. 5b). This shows that LITOS is suitable to screen for perturbations of signaling dynamics with relatively little effort.

Next, we evaluated the potential of LITOS to study ERK dynamics that require high temporal resolution. For this purpose, we stimulated optoFGFR cells with a single 2 second-pulse of blue light and fixed the cells with 1 minute time resolution. This captured highly resolved details of ERK activity dynamics, such as the steep phase of ERK activation followed by a slower phase of adaptation (Fig. 5c). This result emphasizes the ability of LITOS to study rapid signaling events.

The experiments we documented so far are based on single blue-light pulses. However, different signaling networks are known to encode different dynamic signaling patterns that can consist of transient, sustained, oscillatory or pulsatile signaling dynamics^18^. Reproducing these complex dynamics profiles using a synthetic biology approach might provide key insights into the network that shapes these dynamics. Therefore, we evaluated the capability of LITOS to produce more complex light input patterns to evoke specific synthetic ERK dynamics patterns. To do so, we programmed LITOS to stimulate optoFGFR cells with either high (every 2 minutes) or low (every 20 minutes) frequency 10 seconds-long light pulses for a total of 84 minutes. We found that high-frequency light pulses evoke sustained ERK dynamics (Fig. 5d). This is consistent with the finding that activation of optoFGFR by a single blue light pulse switches it on for approximately 5-10 minutes^5^, thus not allowing ERK deactivation upon reactivation every 2 minutes. In contrast, low frequency light pulses evoked successive, discrete ERK pulses that all displayed adaptation to the baseline until a new light input was applied. This shows how LITOS can be used to precisely tune user-defined synthetic ERK dynamics. This experiment required applying different light stimulation schemes to 86 wells of the well plate, illustrating the complexity of the experiments LITOS can tackle. Altogether, these results show that LITOS is a highly flexible optogenetic device that allows one to perform complex experiments to study the dynamics of the MAPK pathway.

## Discussion

We present LITOS, a user-friendly, inexpensive, easy to assemble and robust LED-matrix tool with the aim of democratizing incubator-based optogenetic stimulation experiments. LITOS is composed of parts that can be easily purchased online or ordered from the manufacturer. Here and in a GitHub repository (https://github.com/pertzlab/LITOS), we provide instructions to assemble and use LITOS. We also document a dedicated software solution to control LITOS. We benchmark LITOS by studying the dynamics MAPK signaling pathway, which can be manipulated by different optogenetic actuators and measured using a fluorescent biosensor. This demonstrated that the LED density in LITOS is sufficient to uniformly stimulate all the cells cultured in well-plates (Fig. 4c), enabling to perform reliable population-average measurements using biochemical methods such as western blot (Fig. 4d). Moreover, we showed that LITOS is compatible with two different optogenetic actuators (Fig. 5a), provides the necessary throughput to perform small drug screens (Fig. 5b), allows for high temporal resolution measurements (Fig. 5c) and enables the application of complex light stimulation schemes (Fig. 5d). These features meet the needs of the rapidly expanding field of optogenetics and its application in the study of dynamic processes.

The importance of the dynamic regulation of biological processes, including signaling dynamics, has been increasingly recognized in the last years^18^. The pulsatile dynamics of ERK activity observed in epithelial collectives is one of the most striking examples^16,17,19–22^. Here, the frequency of ERK pulses of constant amplitude and duration has been shown to control different fate decisions, such as cell death, survival, proliferation and migration^16,19,21,22^. Fluorescent biosensors have allowed us to visualize these signaling events at adequate spatial and temporal scales. Optogenetics now also allows us to control these biological processes at these biologically-relevant time and length scales in a non-invasive way. We had previously shown that applying frequency-modulated optoFGFR or optoRaf inputs can evoke frequency-modulated ERK dynamics regimes in which all the cells in the population are responding synchronously^16^. This is in contrast to the commonly used sustained bulk application of growth factors that often lead to heterogeneous, unsynchronized signaling behaviors^16,19,23^. Here, LITOS again reveals that this occurs when specific light-mediated optoFGFR inputs are applied. Application of one light-mediated optoFGFR input with LITOS led to population-homogeneous steep ERK activation followed by adaptation, leading all the cells of a population to experience an identical ERK pulse (Fig. 4c). Multiple light pulses, applied at low frequency, could lead to population synchronous discrete ERK pulses (Fig. 5d). In contrast, when those light pulses were applied at high frequency, that did not allow for optoFGFR inactivation^5^, sustained ERK activity could be evoked (Fig. 5d). We envision that these features of LITOS, as well as of similar devices, will increase the reliability of biochemical population average measurements such as western blot, transcriptomics or (phospho)proteomics. Here, the ability to induce population synchronous ERK dynamics will provide for reliable population average measurements using biochemical approaches. This is in contrast to bulk growth factor stimulation experiments that might average heterogeneous signaling states. For example, we envision that highly temporally resolved phosphoproteomes of an ERK pulse evoked by a transient optoFGFR input might map the fluctuations of phosphosites at biologically relevant timescales, providing a better understanding of receptor tyrosine kinase-MAPK signaling. Alternatively, the ability to evoke desired synthetic ERK dynamics that can induce specific fate decisions (e.g. survival, proliferation, differentiation), when coupled to omics approaches (e.g. transcriptomics, (phospho)proteomics) might provide new insights about how ERK dynamics are decoded into fate decisions. By taking advantage of the ever-expanding repertoire of optogenetic actuators and biosensors, we envision that a similar experimental logic can be applied to other signaling systems.

Another important potential of LITOS is in the study of biological processes that occur over timescales over multiple days that would require dedicated microscopes for large amounts of time for optogenetic stimulation. This is illustrated by a separate study in which we used LITOS to optogenetically control ERK signaling in 3D mammary acini morphogenesis, a developmental process that unfolds over a period of 2 weeks^24^. We focused on an ERK-dependent developmental episode in which the acinar lumen is formed through survival of the outer cells and apoptosis of the inner cell mass in a process that takes approximately one week. We found that the ERK pulse frequency was higher in the outer cells than in the inner cells, and hypothesized that the ERK pulse frequency therefore dictates survival versus apoptosis fates. Using optoFGFR and optoRaf actuators, we used LITOS to evoke different synthetic frequency-modulated ERK pulse regimes. We observed that high frequency ERK pulse regimes rescued the survival phenotypes in inner cells, hampering acinus lumen formation. These results illustrate how LITOS can control morphogenetic events that occur on multiple days in experiments performed within a tissue culture incubator. We envision that LITOS-mediated control of optogenetic actuators could be used for a wide range of applications in 3D biology, including organoid culture.

The simple design of LITOS makes it versatile and affordable. While we restricted our study to CRY2-based systems, LITOS’s RGB LED matrix should also enable control of optogenetic actuators based on cryptochrome, dronpa, and LOV (e.g. iLID, TULIP) domains^25^. However, LITOS’s simplicity might also come with some limitations. Its light intensity is lower than the one produced by a typical fluorescence microscope. We recommend new users to test if their favorite optogenetic system can be activated by the light intensity produced by LITOS. Moreover, LITOS is not spectrally compatible with phytochrome-based actuators that require infrared light.

In sum, we developed and tested LITOS, a cheap, robust and easy to use LED illumination device for optogenetics. We have illustrated how LITOS could be used in the study of MAPK signaling. We envision a broad range of applications for LITOS outside the signaling field. For example, it could be applied to study light-controlled gene expression in bacteria^26^, developmental biology research in Drosophila^27^ or for CRISPR-Cas9 photoactivatable genome editing^28^.

Further information about LITOS, how to order one, and its open source code are available in the following GitHub repository (https://github.com/pertzlab/LITOS).

## Material and Methods

### LITOS components

LITOS consists of a commercially available RGB LED matrix (P3 indoor LED display module, 32×64 Pixel, with FM6126A IC, from Shenzhen Xuyang Technology Co. ltd., albeit modules with other ICs could be potentially used as well) for the cell stimulation and a custom printed circuit board (PCB). The peak emissions of the LED of the RGB matrix is 620-630 nm for red, 520-525 nm for green and 465-470 nm for blue. The PCB is built around the ESP32-WROOM module (Espressif systems Shanghai Co. ltd.) and acts as a control unit for the LED matrix. Besides the circuits required for this task, an 1.5-inch OLED screen module (with SSD1351 control IC), control buttons (B3F-4050, Omron) and a buzzer (AI-1223-TWT-3V-2-R, PUI audio) provides additional audiovisual input/output possibilities. Initial programming can be conducted by the added USB serial UART bridge (231-XS, FTDI). Power is supplied by an external 5-volt wall power adapter, connected to a 2.1 mm x 5.5 mm barrel jack connector placed on the PCB. As there are components requiring a lower voltage of 3.3V, a linear voltage regulator (LM1117, Texas Instruments) was added to the PCB. LITOS’s circuit diagram, the PCB layout files (generated with KiCad 6.01, https://www.kicad.org/) and its bill of material can be found in our GitHub repository (https://github.com/pertzlab/LITOS).

### LITOS assembly

The PCB is connected to the RGB LED matrix with a cable that provides the required power and with a 16-pin flat ribbon cable used for signal transmission. Both cables connect to the PCB and to the RGB LED matrix through existing headers. Finally, the PCB is connected to the external 5V power supply. Furthermore, the PCB can be connected to a computer via a USB cable. This is required to initially load LITOS’ software onto the ESP32 microcontroller (if not loaded by the PCB manufacturer). After that, the connection to a computer is established over Wi-Fi.

### LITOS control software

The ESP32, located on the PCB, is programmed in C / C++ using the Arduino hardware abstraction layer. The configuration interface is programmed in Vue, a progressive Web framework for building the Configuration Interface as single-page applications (https://vuejs.org/). The source code can be found in our GitHub repository (https://github.com/pertzlab/LITOS).

### Cell culture

We used the mouse embryonic fibroblast cell line NIH3T3 and the non-tumorigenic human mammary epithelial cell line MCF10A. We maintained NIH3T3 cells in Dulbecco’s Modified Eagle’s Medium - high glucose (DMEM) from Sigma-Aldrich (Ref: D5671) supplemented with 10 % fetal bovine serum (FBS), 200 U/ml penicillin and 200 µg/ml streptomycin and 200 nM L-glutamine. Before passaging, the confluency was kept below 40-50 % to avoid changes in the behavior of NIH3T3 cells. The MCF10A were maintained in DMEM:F12 (D8437) from Sigma-Aldrich supplemented with 5% Horse Serum, 200 U/ml penicillin and 200 µg/ml streptomycin, 20 ng/ml EGF (Peprotech), 10 µg/ml Insulin (Sigma-Aldrich/Merck) and 0.5 µg/ml of Hydrocortisone (Sigma-Aldrich/Merck).

NIH3T3 and MCF10A cells were starved in a medium consisting of Ham’s F-12 Nutrient Mixture (Sigma N4888) with 200 U/ml penicillin and 200 µg/ml streptomycin and 200 nM L-Glutamine and 0.5 % bovine serum albumin for the NIH3T3 cells and DMEM:F12 (Sigma D8437) supplemented with 0.3% BSA, 0.5 µg/ml Hydrocortisone and 200 U/ml penicillin and 200 µg/ml streptomycin for MCF10A cells. NIH3T3 and MCF10A cells were stably modified to express optoFGFR (fused with a mCitrine fluorescent protein) or optoRAF (fused with a mCitrine fluorescence protein as well). OptoFGFR was transduced by a lentiviral vector (Lyn-cytoFGFR1-PHR-mCit) and optoRAF by a PiggyBac plasmid (pPB3.0.PURO.CRY2.cRAF.mCitrine.P2A.CIBN.KrasCT). Both the systems express puromycin resistance that we used to select stably transduced cells. The cell lines expressing these optogenetic actuators were further transfected with two additional PiggyBac plasmids to stably express ERK-KTR-mRuby2 with hygromycin resistance and the nuclear marker H2B-miRFP703 with blasticidin resistance. For NIH3T3 cells, the PiggyBac plasmids were transfected using jetPEI transfection reagent. For the MCF10A cells, the plasmids were transfected with the FuGENE HD Transfection Reagent (Sigma). After transfection, the cells were selected with antibiotics. Additionally, NIH3T3 were FACS sorted while MCF10A cells were clonally expanded to increase homogeneity of expression of the biosensors.

### LITOS optogenetic experiments

We seeded 2 ∗ 10^³^ NIH3T3 cells / well in a 96 well plate for the imaging experiments and 1.5 * 10^6^ MCF10A cells / well in a 6 well plate for western blotting. In both the cases, we used optical glass-bottom multi-well plates coated with 10 μg/mL fibronectin for one hour before seeding. The cells were starved for 24 h prior to the start of the experiment.

For all experiments, the LITOS illumination patterns were created in LibreOffice Calc. To upload those files, we connected a laptop to LITOS’s access point, and used the user interface. Once the illumination patterns were uploaded on LITOS, the multi-well plate was placed on the LED matrix in the incubator. To ensure proper placement of the well plate, we used a plexiglass mask purpose-made with a CNC milling machine (Fig. S4). Cells were allowed to adapt for at least one hour to the incubator’s conditions before an optogenetic experiment was started. If allowed by the experimental protocol, we avoided opening the incubator during the experiment. After stimulation, cells were fixed by adding the same volume of 4 % paraformaldehyde solution to the volume of medium present in the well. For all shown experiments, the different experimental points were stimulated at the specific time points to be then fixed simultaneously at the end of the experiment. After a 10-min of fixation, we washed the cells twice with PBS.

### Cell imaging and single-cell ERK activity measurements

Widefield fluorescence microscopy images were acquired using a Nikon Eclipse Ti Microscope equipped with 20x Plan Apo Lambda (NA 0.75) and an Andor Zyla 4.2+ camera. For excitation, we used the following LED light sources: 555 nm for ERK-KTR and 640 nm for nuclear marker). For stimulation of optoFGFR we used a 488 nm LED light source. Imaging analysis was done with a pipeline established in our laboratory^16^. First, using the H2B nuclear marker, we created a probability map of the nuclei with a random forest classifier based on different features using Ilastik software (version 1.3.3). Based on this probability map, we used CellProfiler (version 3.0.0) to segment the nucleus. We then segmented the cytosol in a ring shape starting 2 px away from the nucleus and a maximum width of 5 px (Fig. 4b). The 2 px distance between the nucleus and cytosol segmentation ensures that imprecision in nucleus segmentation does not impact the results. After measuring the ERK-KTR signal with those two segmentation masks, the ratio between the ERK-KTR present in the cytosol and in the nucleus (C/N-ratio) was calculated. The relative cytoplasmic versus nuclear fluorescence of the KTR construct (C/N ratio) correlates with the ERK activity and can be used as a proxy for the kinase activity in single cells.

### Western blot

For western blotting, cells were first fixed with 4 % paraformaldehyde for 10 minutes and then lysed in 2 % SDS buffer (in TrisHCl pH 6.8). Protein concentration was determined with Pierce BCA Protein Assay Kit (Ref. 23227, Thermo Scientific). From each cell lysate 10 µg of protein were loaded to an SDS-Page (10 % self-made gel) and then blotted to a PVDF western blot membrane (Ref: 3010040001, Roche). The membrane was incubated with rabbit monoclonal antibody against total ERK (Ref: #4695) or rabbit monoclonal antibody against phospho-ERK (Ref: #4370) both from Cell Signaling, and then with Amersham ECL Rabbit IgG, HRP-linked whole Ab (from donkey, NA934VS), as secondary antibody. The membrane was then developed using Amersham ECL Prime Western Blotting Detection Reagents (RPN2232) from GE Healthcare.

### Statistical analysis

Single cell data are represented with box plots or mean with standard error of the mean, as described in the relative figure legend. Data analysis and figures were created with R (version 3.6.3, https://www.R-project.org/)^29^. Corresponding data and R-notebook files used for data analysis and image creation can be found in the Supplementary Material.

## Supporting information

Data and R-scripts

## Acknowledgements

This work was supported by a grant from the Swiss National Science Foundation to Olivier Pertz. We are grateful to the technical workshop of the Department of Chemistry, Biochemistry and Pharmaceutical Sciences and Microscopy Imaging Center (https://www.mic.unibe.ch) of the University of Bern for technical support.

## Author contributions

T.H. conceived the optogenetic plate design. A.L. did the hardware and software development. C.D. built the genetic circuit consisting of optoFGFR, optoRaf and the ERK-KTR. P.A.G. produced the MCF10A cell lines with the optoFGFR, optoRaf and ERK-KTR. T.H. conducted and analyzed cell based experiments. T.H. created all figures. O.P. supervised the work. T.H., P.A.G., and O.P. wrote the paper. All authors reviewed the manuscript.

## Data availability statement

All datasets, including the R notebooks used for analysis, are included in the supplementary information.

## Additional Information

The authors declare no competing interests.

