## Supplementary figures and legends for "LITOS - a versatile LED illumination tool for optogenetic stimulation"

**Supplement Figure S1**

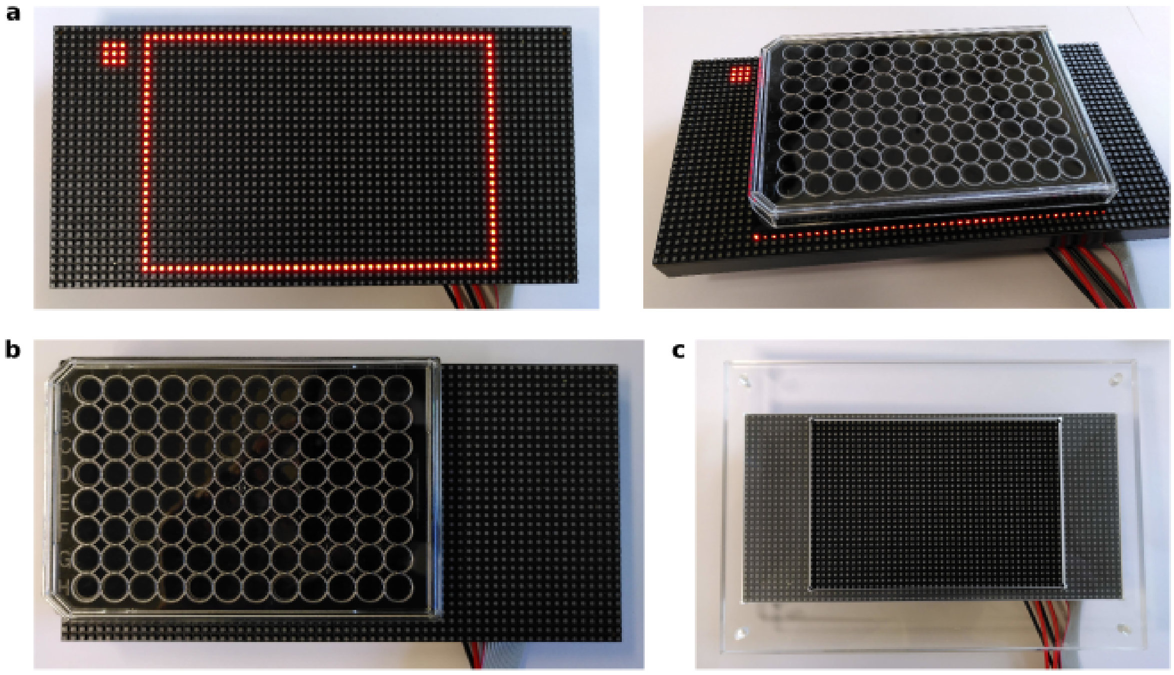


**Figure S1: Different possibilities to align a well plate on the LED matrix.** a) A way to align (especially in the dark) a multi-well plate on the LED matrix of LITOS is to use LITOS to show a reference outline on the matrix. The 3 by 3 illuminated square indicates the upper left-corner of the multi-well plate. b) Alignment of the plate to the top left corner of the LED matrix. c) A plexiglass mask is used to position one multi-well plate on the LED matrix (the plexiglass mask to position two multi-well plates is shown in figure 2C). The position of the illumination patterns can be set in the configuration interface according to the alignment approach.

**Supplement Figure S2**

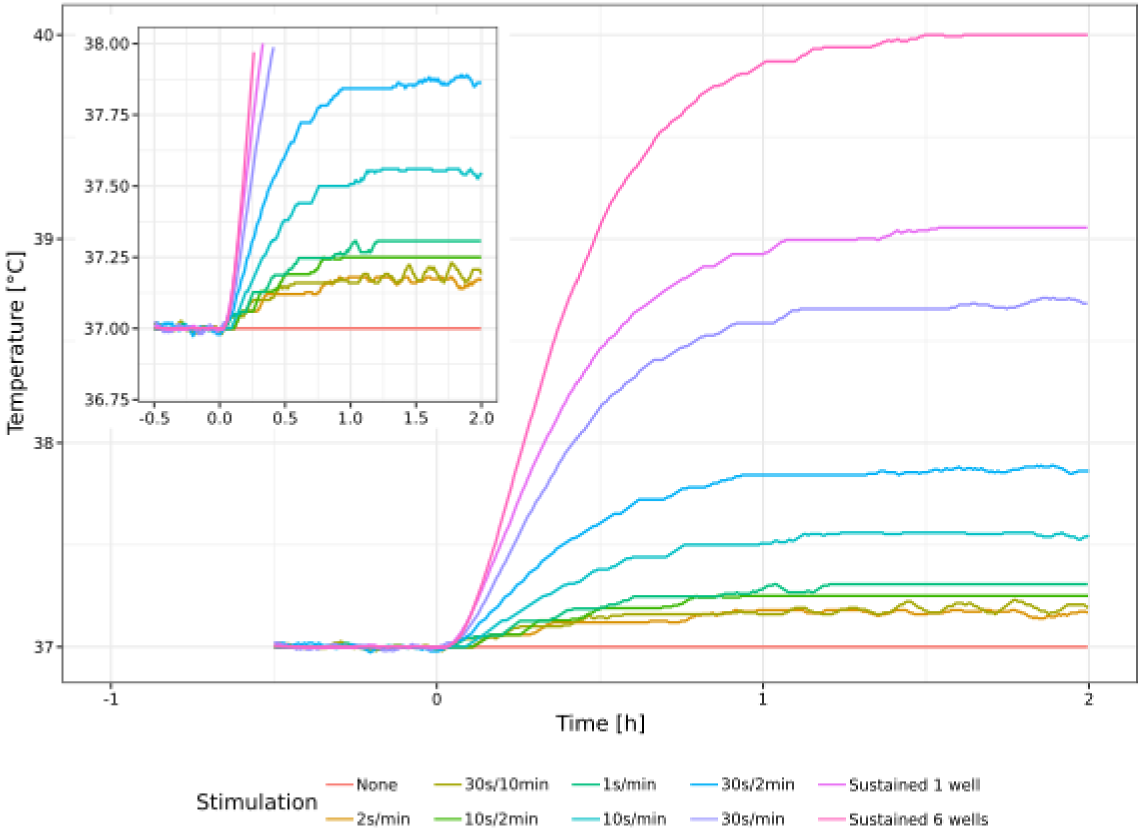


**Figure S2: Increase in medium temperature due to different illumination frequencies.** The temperature increase in an incubator due to different illumination intervals was determined by using an Arduino device equipped with multiple DS18B20 waterproof temperature sensors. The temperature sensors were placed in a well (6 well plate) filled with 3 ml water.

**Supplement Figure S3**

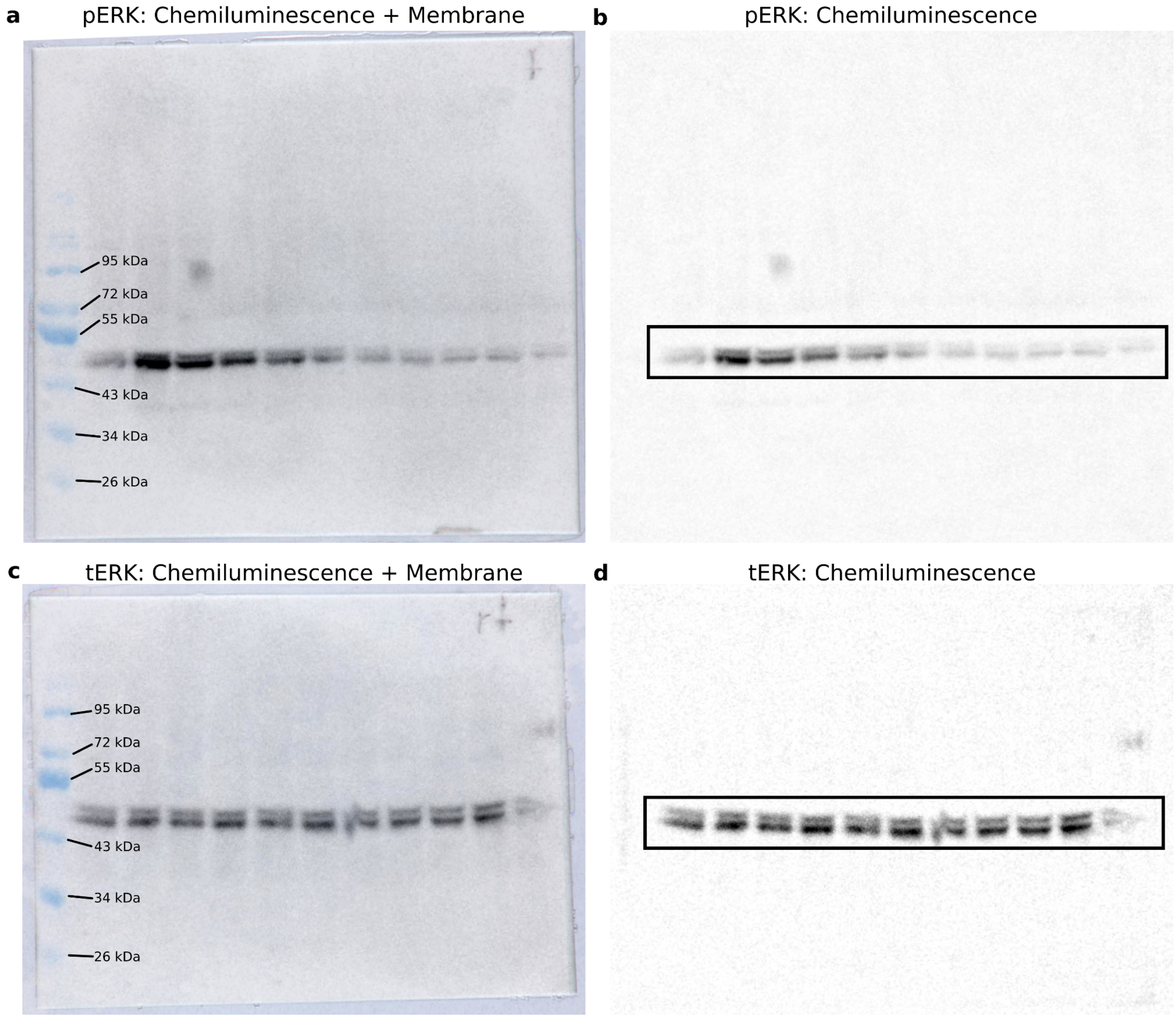


**Figure S2: Uncropped plots from Figure 4d.** Western blot against phosphorylated (a, b) and total ERK (c, d). Time series from 0 min to 27 min after optogenetic stimulation, with 3 min resolution.

**Movie S1**

Movie S1: Movie showing a stimulation pattern generated by LITOS for 96 well plates, in which every column is stimulated sequentially. Illumination with blue light at maximum intensity.
