## Supplementary material for "LITOS - a versatile LED illumination tool for optogenetic stimulation": Data and R-scripts: 1_cell_based_experiemnts.nb.html

Cell based experiments to showcase LITOS


Code 

- Show All Code
- Hide All Code
- Download Rmd

### Cell based experiments to showcase LITOS

### Introduction

This is the data analysis code of the cell-based experiment (Figure 4 and 5) to showcase LITOS with the optogenetic stimulation of the MAPK/ERK pathway. The methodological details about the experiments and the description of the results can be found in the main text of the paper.

### Analyses of the experiments

#### Loading packages and data


The meta-package `tidyverse` is used to prepare (e.g. with `dplyr`) and to plot (with `ggplot2`) the data.

The first step is to load the data.


```
data_western_blot <- read_csv("data_WesternBlot.csv")
data_dose_response <- read_csv("data_DoseResponse.csv")
data_dynamics <- read_csv("data_Dynamics.csv")
data_high_resolution <- read_csv("data_HighResolution.csv")
data_uniform_activation <- read_csv("data_Uniform.csv")
data_mini_screen <- read_csv("data_MiniScreen.csv")
```

#### Uniform activation experiment

Plot the color-coded ERK activity (C/N - ratio) for each cell at its respective position in the well (Figure 4c).


```
data_plot <- data_uniform_activation %>%
  # Manually remove outliers
  filter(!(Image_Metadata_Well == "D09" & objNuc_ObjectNumber == 1) &
           !(Image_Metadata_Well == "D11" & objNuc_ObjectNumber == 207) & 
           !(Image_Metadata_Well == "D11" & objNuc_ObjectNumber == 2885)) %>% 
  
  # Rename columns
  mutate(X = objNuc_Location_Center_X, Y = objNuc_Location_Center_Y, 
         `C/N` = cytoVnuc.corr, 
         Timepoint = paste("t = ", Timepoint, "'", sep = ""), 
         Timepoint = factor(Timepoint, levels = c("t = 0'", "t = 3'", "t = 6'", "t = 9'", "t = 12'"))) %>% 
  
  # Limit the C/N to a min of 0.2 and a maximum of 1.5, so the color scale is more meaningful
  mutate(
    `C/N` = ifelse(`C/N` < 0.2, 0.2, `C/N`), 
    `C/N` = ifelse(`C/N` > 1.5, 1.5, `C/N`
  )) %>%
  ungroup()
          
# Plot the data^
ggplot(data_plot, aes(x = X, y = Y, col = `C/N`)) +
  geom_point(alpha = 0.85, shape=16, size =0.5) +
  scale_y_reverse() + 
  scale_color_gradient(high = "#f75555", low  = "#56B1F7") +
  theme_void() + theme(legend.position = "top") +
  facet_grid(. ~ Timepoint)
```

#### Western blot experiment

Plot the color-coded ERK activity (C/N - ratio) for each timepoint so that they can be compared with the western blot results (Figure 4d).


```
data_plot <- data_western_blot %>% 
  group_by(RealTime) %>%
  summarise(meanResponse = mean(Response), medianResponse = median(Response))

ggplot(data_plot, aes(x = RealTime, y = 1, fill = meanResponse)) + 
  geom_tile() +
  scale_fill_gradient(high = "#f75555", low  = "#56B1F7") +
  theme(legend.position = "bottom") +
  xlab(xlab) + ylab(ylab)
```

#### Dose-response experiment

Plot the dose response experiment (Figure 5a).


```
# Prepare data for plot
data_plot <- data_dose_response %>% 
  mutate(RealTime = factor(RealTime)) # change RealTime to a categorical variable

# Create data for blue square indicating stimulation
data_stim <- tibble(
  Cells = c("optoFGFR1", "optoFGFR1", "optoFGFR1", "optoFGFR1", "optoRAF", "optoRAF", "optoRAF", "optoRAF"),
  ExposureTime = c("0.5 min", "1 min", "2 min", "5 min", "0.5 min", "1 min", "2 min", "5 min"), 
  start = 0,
  end = c(0.5, 1, 2, 5, 0.5, 1, 2, 5),
  min = 0.2,
  max = 1.5
  ) 

# Plot 
ggplot(data_plot) +
  geom_boxplot(aes(x = RealTime, y = Response), outlier.shape = NA) +
  geom_rect(data=data_stim, aes(xmin = start+1, xmax = (end/3)+1, ymin = min, ymax = max), fill = "blue", alpha = 0.2) +
  geom_boxplot(aes(x = RealTime, y = Response), outlier.shape = NA) + # put boxplot over rectangles
  coord_cartesian(ylim = c(0.27, 1.3)) +
  facet_grid(Cells ~ ExposureTime) +
  theme_bw() + theme(legend.position = "bottom", text = element_text(family = "sans-serif", size = 8)) +
  xlab(xlab) + ylab(ylab)
```

#### Mini drug screen

Plot the results of the mini drug screen which was performed with LITOS (Figure 5b).


```
# Calculate mean per time point
data_plot <- data_mini_screen %>%
  group_by(Drug, Concentration, RealTime) %>% 
  summarise(err = sd(Response)/sqrt(n()), Response = median(Response), .groups = "drop")

# Generate data to indicate stimulations
data_stim <- tibble(
  Drug = c("RAF709", "UO126", "SCH772984")
  ) 

ggplot(data_plot) +
  geom_rect(data = data_stim, aes(xmin = 0.3, xmax = 0.7, ymin = 0.2, ymax = 0.7), fill = "blue", alpha = 0.2) +
  geom_point(aes(x = RealTime, y = Response, col = Concentration)) +
  geom_line(aes(x = RealTime, y = Response, col = Concentration, group = paste(Concentration, Drug))) +
  coord_cartesian(ylim = c(0.44, 0.65)) +
  theme(text = element_text(family = "sans-serif", size = 8), legend.position = "top") +
  facet_wrap(~Drug) +
  xlab(xlab) + ylab(ylab)
```

#### High temporal resolution experiment

Plot a single ERK activity pulse with high temporal resolution (Figure 5c).


```
data_plot <- data_high_resolution

# Generate data to indicate stimulations
dp_stim <- tibble(
  Drug = c("RAF709")
  ) 

ggplot(data_plot) + 
  geom_boxplot(aes(x = factor(Timepoint), y = cytoVnuc.corr), outlier.shape = NA) + 
  geom_rect(data=dp_stim, aes(xmin = 1.3, xmax = 1.5, ymin = 0, ymax = 1.6), fill = "blue", alpha = 0.2) +
  geom_boxplot(aes(x = factor(Timepoint), y = cytoVnuc.corr), outlier.shape = NA) +  # put boxplot over rectangles
  coord_cartesian(ylim = c(0.15, 1.438)) +
  theme(text = element_text(family = "sans-serif", size = 8)) +
  xlab(xlab) + ylab(ylab)
```

#### Induction of temporal dynamics

Plot the dynamics which was generated with LITOS (Figure 5d).


```
# Calculate mean and SEM per time point
data_plot <- data_dynamics %>% 
  # mutate(RealTime = as.numeric(as.character(RealTime))) %>%
  group_by(RealTime, PulseInterval) %>%
  summarise(mean = mean(Response), sd = sd(Response), sem = sd(Response)/sqrt(n()), .groups = "drop")

# Generate data to indicate stimulations
min = min(data_plot$mean)
max = max(data_plot$mean)

data_stim <- tibble(
  PulseInterval = c("sustained", "every 20 min", "every 20 min", "every 20 min", "every 20 min"),
  start = c(4, 4, 24, 44 , 64),
  end = c(84, 4.4, 24.4, 44.4, 64.4),
  min = min-0.1,
  max = max+0.1
  ) 

# Plot 
ggplot(data_plot) +
  geom_rect(data=data_stim, aes(xmin = start, xmax = end, ymin = min, ymax = max), fill = "blue", alpha = 0.2) +
  geom_line(aes(x = RealTime, y = mean, group = PulseInterval)) +
  geom_errorbar(aes(x = RealTime, y = mean, ymin = mean - sem, ymax = mean + sem)) +
  facet_grid(PulseInterval ~ .) +
  theme(text = element_text(family = "sans-serif", size = 8)) +
  xlab(xlab) + ylab(ylab)
```
