## Supplementary material for "LITOS - a versatile LED illumination tool for optogenetic stimulation": Data and R-scripts: 2_temperature_measurement.nb.html

Temperature increased caused by LITOS


Code 

- Show All Code
- Hide All Code
- Download Rmd

### Temperature increased caused by LITOS

### Experiment

LITOS can warm up when it is used. This is especially the case, when it is used for prolonged sustained stimulations with optogentics. With shorter and less frequent stimulation pulses the heat development is less a concern. To quantify the warm up of LITOS we used an Arduino device equipped with five DS18B20 waterproof temperature sensors.

Placement of the 5 sensors:

- Incubator: Measure changes in the air temperature of the incubator.
- Negative: Multiwell plate in the incubator as negative control.
- LITOS 1: Whole multiwell plate under sustained stimulation (maximal temperature increase)
- LITOS 2: Multiwell plate on a second LITOS to test different stimulation patterns
- LITOS 3: Multiwell plate on a third LITOS to test different stimulation patterns

The temperature sensors were placed in a well filled with 3 ml water of a six well plate.

Stimulations pattern tested in LITOS 2:

1. 1 s / min (2 h)
2. wait (3 h)
3. 10 s / 2 min (2 h)
4. wait (3 h)
5. 10 s / min (2 h)
6. wait (3 h)
7. 30 s / min (2 h)

Stimulations pattern tested in LITOS 3:

1. 2 s / min (2 h)
2. wait (3 h)
3. 30 s / 10 min (2 h)
4. wait (3 h)
5. 30 s / 2 min (2 h)
6. wait (3 h)
7. sustained one well (2 h)

The waiting time between the different stimulation schemes, ensures that the plate has time to cool down again.

### Analysis

#### Load packages


```
library(tidyverse) # data preparation and plotting
library(TTR) # rolling average

# Settings for plots
theme_set(theme_bw())
n.plot.height = 6
n.plot.width = 8
```


The meta-package `tidyverse` is used to prepare (e.g. with `dplyr`) and to plot (with `ggplot2`) the data.

#### Load and prepare the data

Load the raw data with the following columns:

- time: Time in (h)
- sensor: Used sensor
- temperature: temperature measured by the sensor

Since temperature sensors are better suited to measure changes than absolute values, we use the data before the experiment to normalize all sensors to 37°C. *Note:* 37°C was arbitrary chosen by us, since it is the temperature you would expect in the incubator.


```
raw_data <- read_csv("data_temperature_measurement.csv")

data_normed <- raw_data %>%  
  group_by(sensor) %>% 
  mutate(temperature = temperature - mean(temperature[1:450]) + 37) # normalize all sensor to 37°C
```

#### Check the temperature over the course of the experiment.

In the following plot the measured temperature (change) for each sensor is showed. To smooth out the values a rolling average was used.


```
data_plot <- data_normed %>% 
  group_by(sensor) %>%
  mutate(avg = runMean(temperature, 9)) %>% #take the rolling average
  na.omit() # remove na which where introduced by taking the rolling average

ggplot(data_plot, aes(x = time, y = avg, col = sensor)) +
  geom_line()
```

#### Check temperature changes for individual stimulation patterns

To compare the effect of the individual illumination patterns, we extract them from the data and plot them. We again use a rolling average to smooth the data.


```
data_smooth <- data_normed %>% 
  group_by(sensor) %>%
  mutate(avg = runMean(temperature, 9)) %>% #take the rolling average
  na.omit() # remove na which where introduced by taking the rolling average

# Extract individual "parts" of the experiment
first <- data_smooth %>% 
  mutate(part = "first", 
         time = time) %>%
  filter(time > -0.5, time < 2)

second <- data_smooth %>% 
  mutate(part = "second", 
         time = time - 5) %>%
  filter(time > -0.5, time < 2)

third <- data_smooth %>% 
  mutate(part = "third", 
         time = time - 10) %>%
  filter(time > -0.5, time < 2)

fourth <- data_smooth %>% 
  mutate(part = "fourth", 
         time = time - 15) %>%
  filter(time > -0.5, time < 2)

# Combine the parts together
data_plot <- union(first, second) %>% 
  union(third) %>% 
  union(fourth) %>% 
  # Normalize the part bevor the stimulation pattern to 73°C
  group_by(sensor, part) %>% 
  mutate(avg = avg - mean(avg[1:85]) + 37) %>% # take rolling average
  na.omit() %>% # remove NA introduced by rolling average
  # Name the stimulations pattern 
  mutate(Stimulation = ifelse(sensor == "Negative" & part == "first", "None", 
                              ifelse(sensor == "Sustained" & part == "first", "Sustained 6 wells", 
                                     ifelse(sensor == "LITOS2" & part == "first", "1s/min", 
                                           ifelse(sensor == "LITOS2" & part == "second", "10s/2min", 
                                                  ifelse(sensor == "LITOS2" & part == "third", "10s/min", 
                                                         ifelse(sensor == "LITOS2" & part == "fourth", "30s/min", 
                                                                ifelse(sensor == "LITOS3" & part == "first", "2s/min", 
                                                                       ifelse(sensor == "LITOS3" & part == "second", "30s/10min", 
                                                                              ifelse(sensor == "LITOS3" & part == "third", "30s/2min", 
                                                                                     ifelse(sensor == "LITOS3" & part == "fourth", "Sustained 1 Well", 
                                                                                            ifelse(sensor == "Incubator" & part == "first", "Incubator", NA)))))))))))) %>%
  # Filter for stimulation pattern of interest
  filter(Stimulation %in% c("None", "Sustained 6 wells", "1s/min", "2s/min", "10s/2min", "30s/10min", "10s/min", "30s/2min", "30s/min", "Sustained 1 Well")) 

# Plot the data
ggplot(data_plot, aes(x = time, y = avg, col = Stimulation)) +
  geom_line()+ 
  xlim(-1,2) +
  theme(legend.position = "bottom") +
  xlab("Time [h]") + ylab("Temperature [°C]")
```


```
# Zoomed in plot
ggplot(data_plot, aes(x = time, y = avg, col = Stimulation)) +
  geom_line()+ 
  ylim(36.8,38) + 
  theme(legend.position = "none", text = element_text(size=6)) +
  xlab("") + ylab("")
```
